# Naturally-segregating Variation at *Ugt86Dd* Contributes to Nicotine Resistance in *Drosophila melanogaster*

**DOI:** 10.1101/134429

**Authors:** Chad A. Highfill, Jonathan H. Tran, Samantha K. T. Nguyen, Taylor R. Moldenhauer, Xiaofei Wang, Stuart J. Macdonald

## Abstract

Identifying the causative sequence polymorphisms underlying complex trait variation is a key goal of evolutionary and biomedical genetics. By knowing the precise molecular events that confer phenotypic changes we can gain insight into the pathways underlying complex traits and the evolutionary forces acting on variation. Genetic analysis of complex traits in model systems regularly starts by constructing QTL maps, but generally fails to identify causative nucleotide-level polymorphisms. Previously we mapped a series of QTL contributing to resistance to nicotine in a *Drosophila melanogaster* multiparental mapping resource, and here use a battery of functional tests to resolve QTL to the molecular level. One large-effect QTL resided over a cluster of UDP-glucuronosyltransferases, and quantitative complementation tests using deficiencies eliminating subsets of these detoxification genes revealed allelic variation impacting resistance. RNAseq showed that *Ugt86Dd* had significantly higher expression in genotypes that are more resistant to nicotine, and anterior midgut-specific RNAi of this gene reduced resistance. We discovered a naturally-segregating 22-bp frameshift deletion in *Ugt86Dd*, and overexpression of the insertion-containing allele in a range of tissues enhanced resistance. Accounting for the InDel event during mapping largely eliminates the QTL, implying the InDel explains the bulk of the effect associated with the mapped locus. Finally, we edited a relatively resistant genetic background to generate lesions in *Ugt86Dd* that recapitulate the naturally-occurring putative loss-of-function allele, and succeeded in radically reducing resistance. The putatively causative coding InDel in *Ugt86Dd* can be a launchpad for future mechanistic exploration of xenobiotic detoxification.

**ARTICLE SUMMARY:** Resolving the mutations that contribute to among-individual trait variation is a major goal of biomedical and evolutionary genetics. In general however, genetic mapping experiments do not allow immediate resolution of the underlying causative variants. Previous mapping work revealed several loci contributing to nicotine resistance in *Drosophila melanogaster*. We employed a battery of functional tests to demonstrate that the detoxification gene *Ugt86Dd* has a major phenotypic effect, and that a segregating frameshift mutation is likely causative. Editing the gene to introduce a premature stop codon led to a significant reduction in resistance, validating its role in xenobiotic detoxification.

## INTRODUCTION

A principal goal of research on the genetics of complex, polygenic traits is to identify the precise sequence-level polymorphisms responsible for phenotypic variation. This is either carried out directly in natural populations, or in laboratory mapping panels derived from a sample of naturally-derived chromosomes. The pursuit of causative alleles is open to criticism (Rockman 2012), since with finite power it is necessarily the case that experimentally-identified causative variants represent a biased set of the complete collection of sites impacting variation. Nonetheless, genomewide mapping studies have provided important contributions to our understanding of polygenic trait variation. Robust, replicable associations from unbiased GWAS (genomewide association studies) have provided considerable insight into the pathways underlying disease risk, including identifying novel pathways in the control of disease (Hirschhorn 2009; Visscher *et al.* 2012). This is despite the fact that even massive meta-analytical GWAS miss large numbers of subtle-effect polymorphisms, and localize to specific sites only a small fraction of the heritable variation (Yang *et al.* 2010; Wood *et al.* 2014). In addition, the genes and molecular lesions contributing to an array of crop domestication traits have been identified (Doebley *et al.* 2006; Gross and Olsen 2010), providing detail on the specific molecular differences between modern crop plants and their wild progenitors, and an understanding of the nature of the selection applied.

In model organisms, the hunt for causative, naturally-segregating variation commonly begins with linkage-based QTL (quantitative trait locus) mapping. Whether initiated with two parental strains (Lander and Botstein 1989), or more recently with several founders (Kover *et al.* 2009; Churchill *et al.* 2012; King *et al.* 2012b; Threadgill and Churchill 2012), such mapping designs have tremendous power to find QTL, and in some cases have led to the identification of specific polymorphisms contributing to complex trait variation (e.g., Long *et al.* 2000; Deutschbauer and Davis 2005; Bendesky *et al.* 2011; Cook *et al.* 2016; Linder *et al.* 2016). These variants facilitate a deeper understanding of specific biomedically-relevant traits, and collectively add to a fundamental appreciation of complex trait variation and its maintenance in populations. However, these successes aside, it is quite clear that with the possible exception of yeast - where one can test vast numbers of recombinants, minimizing the statistical challenges associated with finding small-effect variants at high resolution (Ehrenreich *et al.* 2010) - linkage-based genomewide mapping for most complex traits in most systems does not result in the identification of a causative mutation. Even those studies using advanced generation intercrosses (e.g., Parker *et al.* 2014), generally yield insufficient information with which to directly implicate causative variants.

The difficulty with moving from QTL to causative site is compounded if traits are highly polygenic. Cornforth and Long (2003) have shown by simulation that standard QTL mapping approaches are unable to discriminate between a single QTL of large effect, and a series of very closely-linked QTL that each have independent effects. That such collections of adjacent causative sites exist is amply demonstrated by examples where high-resolution analysis succeeds in “fractioning” a single QTL into multiple causative loci (Pasyukova *et al.* 2000; Steinmetz *et al.* 2002; Kroymann and Mitchell-Olds 2005), from elegant work showing that multiple alleles combine to yield the major effect of the *Adh* gene on alcohol dehydrogenase expression in flies (Stam and Laurie 1996), and from recent work in multiparental mapping panels showing that QTL frequently appear to be generated by more than one segregating site (Baud *et al.* 2013; King *et al.* 2014). Thus, to follow up QTL mapping, and identify functional allelic variation at the sequence level, additional molecular and functional tests are typically required.

In Marriage et al. (2014) we described initial genetic dissection of nicotine resistance, specifically the fraction of first-instar *D. melanogaster* larvae emerging as adults on media containing nicotine. We chose nicotine as convenient compound allowing high-throughput phenotyping, facilitating dissection of the genetic control of xenobiotic metabolism and resistance. Using RILs from the DSPR (*Drosophila* Synthetic Population Resource), a set of inbred strains derived from a multiparental, advanced generation intercross (King *et al.* 2012b), we mapped a number of QTL. Two loci - Q1 on chromosome 2L and Q4 and chromosome 3R - were particularly interesting. First, their large effects - Q1 and Q4 explain 8.5% and 50.3% of the broad sense heritability, respectively-suggest that individual causative sequence variants may also contribute relatively large effects, facilitating their identification. Although we note that in the case of Q4 evidence indicated that the overall QTL effect may be derived from multiple genes/alleles (Marriage *et al.* 2014), meaning the effects of causative polymorphisms may be individually modest. Second, each QTL interval harbors genes encoding detoxification enzymes; Q1 encompasses a pair of cytochrome P450 genes, and Q4 harbors a set of ten UDP-glucuronosyltransferase (or UGT) genes. These genes are strong *a priori* candidates to causally contribute to trait variation, and good starting points for the identification of causative alleles.

Our goal in the present study was to provide additional evidence that one or more of the detoxification genes found within QTL intervals directly contributes to variation in resistance, employing fine-mapping, expression profiling, tissue-specific RNAi, overexpression analyses and CRISPR/Cas9 genome editing. Our data point to a single, short 22-bp insertion/deletion event in a coding region of *Ugt86Dd* as a major factor contributing to the previously-mapped Q4 locus.

## MATERIALS AND METHODS

### Larval nicotine resistance assay

The assay used was originally described in Marriage et al. (2014), with the only difference here being that larvae were tested in regular, narrow polystyrene fly vials rather than in wells of 24-well plates. Briefly, parental flies were allowed to lay eggs on a standard cornmeal-yeast-molasses medium containing 0.5% activated charcoal, supplemented with a small amount of active yeast paste to elicit egg laying. Subsequently, first instar larvae were collected and placed either on no-drug media or on media containing 0.18 μl/ml nicotine (N3876, Sigma). No-drug and nicotine media were routinely prepared the day prior to larval collection to minimize variation due to nicotine breakdown. Replicate assay vials contained 30 first instar larvae in nearly all cases (see raw data File S1), and the phenotype for each replicate vial is given as the fraction of first instar larvae that ultimately emerge as adults. For every genotype tested we set up several egg-laying vials, and the mean phenotype is given as the average of multiple replicate assay vials.

All animals were reared and tested at 25°C and 50% relative humidity on a 12 hour light/12 hour dark cycle. Those test animals that were the result of crosses were generated by pairing 10 virgin females with 4-5 males over several replicate vials. Prior to initiating such crosses parental flies were allowed to recover from CO_2_ anesthesia for 24 hours.

### Chromosome substitution

Marriage et al. (2014) identified four autosomal QTL contributing to larval nicotine resistance in the DSPR, and found that for each QTL RILs harboring the A4 founder allele were on average more resistant to nicotine than those harboring the A3 allele (see Figure Three in Marriage *et al.* 2014). To broadly examine the phenotypic effects of the A3 and A4 alleles, and to explore dominance, we created chromosome substitution lines. Specifically, using standard fly crosses with balancers, we generated the genotypes A4/A3; A4; A3 and A3/A4; A3; A4 (chromosomes are listed in the order X/Y; 2; 3). We directly compared these substitution genotypes to inbred strains A3 and A4, and carried out intercrosses among strains. For each genotype we measured the phenotype using two replicate no-drug vials to test for variation in overall viability, and six replicate nicotine-containing vials.

To generate chromosome substitutions the following balancer-containing strains were obtained from the Bloomington *Drosophila* Stock Center (BDSC); The double-balancer strain 8316 (w^1118^; wg^Sp-1^/CyO; sens^Ly-1^/TM6B, Tb^1^), the GFP-marked chromosome 2 balancer strain 5702 (w^1^; sna^Sco^/Cyo, Gal4-Hsp70, UAS-GFP), and the GFP-marked chromosome 3 balancer strain 5704 (w^1^; Sb^1^/TM3, Gal4-Hsp70, UAS-GFP, y^+^ Ser^1^).

### Quantitative complementation tests

To test for the effects of allelic variation between strains A3 (relatively susceptible to nicotine) and A4 (relatively resistant to nicotine) at particular genomic regions and at certain genes, we carried out quantitative complementation tests. Virgin females from A3 and A4 were each crossed to males from a series of mutation-containing lines, along with their co-isogenic wildtype control lines, allowing the production of four types of F_1_ progeny per target mutation (A3/mutation, A3/control, A4/mutation, and A4/control). Phenotypes were based on a total of 2-8 replicate no-drug vials, and 8-18 replicate nicotine-containing vials for each genotype. We employed the following model for analysis: *y* = *μ* + *S* + *D* + (*S* × *D*), where *S* is the effect of founder strain (A3 or A4), *D* is the effect of the mutation (comparing either a deficiency or an insertion against a co-isogenic control stock with the wildtype sequence), and *S* × *D* is the interaction. A significant *S* × *D* term indicates a failure to complement (A3 and A4 functionally differ at the locus under test).

Exelixis (Parks *et al.* 2004) and Bloomington Stock Center, or BSC (Cook *et al.* 2012) deficiencies were obtained from the BDSC, specifically 7497 (w^1118^; Df(2L)Exel6011/CyO), 7957 (w^1118^; Df(3R)Exel7306/TM6B, Tb^1^), 7958 (w^1118^; Df(3R)Exel8152/TM6B, Tb^1^), and 26545 (w^1118^; Df(2L)BSC693/SM6a), along with the co-isogenic control strain 6326 (w^1118^). We additionally obtained the DrosDel (Ryder *et al.* 2004) deficiency 9083 (w^1118^; Df(3R)ED5506/TM6C, cu^1^ Sb^1^) and its co-isogenic control 5905 (also w^1118^ but not necessarily otherwise identical to 6326). Note that all deficiencies are maintained over balancers, and since we must discriminate deficiency-containing and balancer chromosomes in first instar larvae, prior to use we substituted all native balancers with GFP-marked versions derived from BDSC stocks 5702 and 5704 (see above). Finally, we obtained several Minos-based insertional mutants (Metaxakis *et al.* 2005), all of which were generated in the 5905 background; 23530 (w^1118^; Mi{ET1}4Cyp28d1^MB03293^), 23587 (w^1118^; Mi{ET1}Cyp28d2^MB02776^), 24834 (w^1118^; Mi{ET1}Ugt86Dj^MB04890^), and 27861 (w^1118^; Mi{ET1}Ugt86Dh^MB11311^).

### RNAseq to identify nicotine resistance genes

Marriage et al. (2014) carried out RNAseq on first-instar larvae to identify genes differentially expressed between DSPR founder strains A3 and A4, and to find genes induced following nicotine exposure. To complement this experiment we selected a set of six (12) lines with relatively high (low) resistance (File S2) from the set of DSPR pB2 RILs that were previously scored for nicotine resistance (Marriage *et al.* 2014). Strains were raised as described above, 30 first instar larvae from each were placed on no-drug or nicotine-supplemented media for 2 hours, after which larvae were flash frozen in liquid nitrogen and stored at −80°C. Subsequently we isolated RNA from each sample using TRIzol reagent (ThermoFisher Scientific, 15596018), and purified using RNeasy columns (Qiagen, 74104). We then pooled equal amounts of total RNA from each sample to generate four RNA pools (high/nicotine, low/nicotine, high/no-drug, low/no-drug), generated an Illumina TruSeq un-stranded mRNA library from each sample, and combined libraries to sequence over a single HiSeq2500 lane and generate single-end 100-bp reads (KU Genome Sequencing Core). Raw reads are available from the NCBI Sequence Read Archive (Accession number SRP102254). Reads were quality trimmed using scythe (v0.991, github.com/vsbuffalo/scythe) and sickle (v1.200, github.com/najoshi/sickle), assembled to the *D. melanogaster* reference genome (NCBI Build 5.3, ccb.jhu.edu/software/tophat/igenomes.shtml) using TopHat (v2.0.10, Trapnell *et al.* 2009; Kim *et al.* 2013), and significant expression differences between relevant pooled samples were identified using Cufflinks (v.2.2.1, Trapnell *et al.* 2010; Trapnell *et al.* 2013).

### RNAi knockdown

We employed the binary Gal4-UAS RNAi system to knockdown expression of several candidate detoxification genes ubiquitously, and specifically in a number of tissues. We crossed males from each Gal4 driver strain to females containing a UAS-RNAi transgene, or to appropriate control females, and assayed F_1_ Gal4-UAS-RNAi progeny. For each genotype we phenotyped 3-5 replicate no-drug vials and 8-10 replicate nicotine vials.

We obtained UAS-RNAi transgene-carrying stocks from the Vienna *Drosophila* Resource Center (VDRC, Dietzl *et al.* 2007). The “GD” library strains harbor *P*-element derived UAS transgenes, while the “KK” library strains harbor phiC31-based transgenes, where all transgenes reside at the same landing site. Specifically, we use VDRC UAS-RNAi strains 6016 (GD, UAS-RNAi-*Ugt86Dd*), 7870 (GD, UAS-RNAi-*Cyp28d1*), 7868 (GD, UAS-RNAi-*Cyp28d2*), 100353 (KK, UAS-RNAi-*Ugt86Dd*), 102626 (KK, UAS-RNAi-*Cyp28d2*), and 110259 (KK, UAS-RNAi-*Cyp28d1*). We directly compare genotypes containing these transgenes to the appropriate control strains 60000 (GD control strain) and 60100 (KK control strain). In addition, we obtained the Transgenic RNAi Project (TRiP, Perkins *et al.* 2015), strain 53892 (UAS-RNAi-*Cyp28d1*) and its co-isogenic control (36304) from the BDSC.

We drove RNAi in all cells and all timepoints using *Act5C*-Gal4 (derived from BDSC 3954 - y^1^ w*; P{Act5C-GAL4}17bFO1/TM6B, Tb^1^), after first substituting the third chromosome balancer in this strain for the GFP-marked version derived from BDSC 5704 (see above). To drive RNAi in specific groups of malpighian tubule cells we employed a series of strains obtained from Julian Dow and Shireen Davies; c42-Gal4, c710-Gal4, c724-Gal4, uro-Gal4 (Rosay *et al.* 1997; Sozen *et al.* 1997; Terhzaz *et al.* 2010). We obtained the anterior midgut Gal4 driver 1099 from Nicolas Buchon (flygut.epfl.ch). Finally, we obtained a number of strains expressing Gal4 in various parts of the gut from the BDSC, specifically strains 1967 (y^1^ w*; P{GawB}34B), 30828 (w*; P{GawB}Alp4^c232^), 30844 (w*; P{GawB}c601^c601^), and 43656 (w*; P{Scr-GAL4.4}1-3).

### Overexpression of *Ugt86Dd*

To complement our RNAi assays, a gain-of-function analysis was performed for *Ugt86Dd* by creating UAS-*Ugt86Dd* overexpression strains. We PCR amplified the gene from both A3 and A4, and simultaneously added *attB* sites, using the primers 5’-GGGGACAAGTTTGTACAAAAAAGCAGGCTTACAACATGAGATTATTAACTGTGATCGCGA-3’ and 5’-GGGGACCACTTTGTACAAGAAAGCTGGGTCCTAATGTTTCTTAAGCTTATCAG-3’. Following the manufacturer’s protocol (ThermoFisher, PCR Cloning System with Gateway Technology, 12535029) we cloned the PCR products into the pDONR221 vector, creating entry clones via BP recombination reactions. Clones were Sanger sequenced using M13 primers to verify the sequence and direction of the insert. The destination vector pUASg.attB was generously donated by Johannes Bischof (Bischof *et al.* 2013) and used in combination with the LR reaction (ThermoFisher, Gateway LR Clonase, 11791020) to generate both pUASg.Ugt86Dd-A3.attB and pUASg.Ugt86Dd-A4.attB expression clones. Expression clones were Sanger sequenced using primer hsp-GW-F (5’-GCAACTACTGAAATCTGCCAAG-3’) to verify the direction of the insert. To generate fly stocks we injected BDSC stock 24749 (y^1^ M{vas-int.Dm}ZH-2A w*; M{3xP3-RFP.attP}ZH-86Fb) with the expression plasmids at 0.510-0.515μg/μl (BestGene, Inc.). We obtained one successful A3 transformant line (UAS-*Ugt86Dd*^A3^), and five A4 transformants (UAS-*Ugt86Dd*^A4(1)^ to UAS-*Ugt86Dd*^A4(5)^), that are each homozygous for the transgene-containing chromosome 3. These strains were utilized in conjunction with tissue-specific Gal4 drivers (above) to explore the effects of *Ugt86Dd* overexpression. For each genotype we tested 2-3 replicate no-drug vials and 6 replicate nicotine vials.

### PCR genotyping assay for *Ugt86Dd* InDel variant

We identified a 22-bp insertion/deletion (InDel) variant in a protein-coding region of exon two in *Ugt86Dd*, and developed a PCR/gel electrophoresis assay to genotype this variant. Primers 5’-CGCTTTTGCTCAGCATTTTA-3’ and 5’-ATATGTGGCAGGTGAACGAA-3’ will amplify 219bp and 197bp products in the presence of the insertion and deletion allele, respectively. PCR cycling conditions were 95°C for 2 min, 35 cycles of 95°C for 20 sec, 57°C for 25 sec, and 72°C for 30 sec, with a final 2 min extension at 72°C. Products were sized on 2% gels run at low voltage for up to 2 hours.

### QTL mapping

Marriage et al. (2014) used the DSPR to identify a highly-significant nicotine resistance QTL encompassing *Ugt86Dd*, along with nine other UGT genes. To test whether the 22-bp InDel in *Ugt86Dd* might contribute to the QTL effect, we first assessed the allelic status of the DSPR founder lines via PCR, finding that founders A3, AB8, B6, and B7 possess the deletion allele (Figure S1). Next, we employed the estimated founder genotype probabilities for the DSPR RILs (King *et al.* 2012b) and determined the allelic status of each RIL at the InDel (File S3). Finally, we mapped nicotine resistance in both pA and pB panels of DSPR RILs following Marriage et al. (2014), but additionally included InDel status as a covariate.

Methods for QTL mapping in the DSPR are described in King et al. (King *et al.* 2012a; King *et al.* 2012b). We employed the R (r-project.org) packages *DSPRqtl*, *DSPRqtlDataA*, and *DSPRqtlDataB* for mapping (FlyRILs.org), analyzed the pA and pB panels of RILs separately, and - in addition to the InDel covariate - included a covariate accounting for the subpopulation from which the RIL was derived (pA1, pA2, pB1, pB2). Note that in the analyses conducted for this study we only employed RILs for which we had high confidence in the founder haplotype at the *Ugt86Dd* gene, so sample sizes are marginally reduced compared to those employed in Marriage et al. (2014), and estimates of the QTL effect are slightly different than previously-reported. To establish significance of mapped QTL we generated genomewide 5% significance thresholds via 1000 permutations of the data (Churchill and Doerge 1994).

### Testing the effect of the InDel variant in the DGRP

We showed via our PCR assay that seven strains from the *Drosophila* Genetic Reference Panel, DGRP (Mackay *et al.* 2012; Huang *et al.* 2014) were homozygous for the deletion allele (BDSC numbers 25176, 25177, 25206, 28185, 28199, 28213, 37525). We also confirmed that the following seven strains were homozygous for the much more common insertion allele (25174, 25200, 28160, 28197, 28226, 28239, 28295). We created a pair of populations, one fixed for the deletion and one fixed for the insertion, by carrying out all possible crosses between the seven founding strains (10 virgin females × 10 males), collecting 10 F_1_ progeny of each sex from each cross, and combining into a 1-gallon population bottle. Both populations were maintained for seven generations, at which point we phenotyped each population using 20 replicate nicotine-containing vials.

### *Ugt86Dd* CRISPR/Cas9 editing

We used the CRISPR Optimal Target Finder (tools.flycrispr.molbio.wisc.edu/targetFinder) to identify an appropriate guide RNA (gRNA) target sequence within *Ugt86Dd* (Gratz *et al.* 2014). Specifically we wished to target the site of the 22-bp InDel, and generate mutations within the *Ugt86Dd* gene of strain A4 that contains the insertion allele. The target sequence 5’-TCACTACGAAGTCATTGTGGAGG-3’ matches the sequence of A4, and was used to generate a gRNA that leads to double-strand breaks within the InDel sequence (Figure S1).

We purchased the 5’-phosphorylated sense and antisense oligos 5’-CTTCGTCACTACGAAGTCATTGTGG-3’ and 5’-AAACCCACAATGACTTCGTAGTGAC-3’, annealed, and followed the protocol from Gratz et al. (2013) to clone the annealed oligos into the pU6-BbsI-chiRNA plasmid (Addgene, plasmid 45946). Transformants were verified via Sanger sequencing.

Our ultimate goal was to compare the phenotypes of inbred strains differing for wildtype and mutant *Ugt86Dd* alleles in an otherwise genetically-identical background. Preliminary sequencing revealed that chromosome 3 for the *vasa*-Cas9 BDSC strain 55821 (y^1^ M{vas-Cas9.RFP-}ZH-2A w^1118^) is not homozygous. Thus, we used a series of standard fly crosses with balancers to substitute chromosome 3 from inbred strain A4 into 55821, creating a strain homozygous for *vasa*-Cas9 on the X and A4 on chromosome 3 (the state of chromosome 2 is unknown, but is likely heterozygous).

The gRNA plasmid was injected into 300 embryos from our custom *vasa*-Cas9; A4 injection strain at 250ng/μl (BestGene, Inc.) yielding 89 G_0_ progeny, 57 of which were shown to be fertile after being individually crossed to 1-2 animals from BDSC 5704 (see above). Where possible we collected 10 F_1_ balanced progeny from each G_0_ cross, and individually crossed to 1-2 animals from BDSC 5704. For crosses containing a mutation (see below), and for some that did not, we collected balanced F_2_ progeny of both sexes to establish stocks, removing the balancer in subsequent generations to create strains carrying homozygous third chromosomes. Note that these strains can and do segregate on the X and chromosome 2.

After each F_1_ cross produced eggs, we genotyped the F_1_ animal for the presence of a CRISPR/Cas9-induced mutation using T7 endonuclease (NEB, M0302L). Briefly, following single-fly DNA isolation (Qiagen, Puregene Cell and Tissue Kit, 158388) we PCR amplified a region surrounding the gRNA target site using oligos 5’-ACGCTTTTGCTCAGCATTTT-3’ and 5’-GGCTGGGGATACCATTTCTT-3’ (95°C for 2 min, 35 cycles of 95°C for 20 sec, 57°C for 25 sec, and 72°C for 30 sec, with a final 2 min extension at 72°C). Subsequently 10μl of PCR product was mixed with 7μl of molecular grade water and 2μl of NEB buffer 2 (B7002S), incubated at 95°C for 5 min in a heat block, and allowed to slowly cool to room temperature for ∼2 hours. We then added 1μl of T7 endonuclease to each reaction, incubated at 37°C for 15 min, added 2 μl of 0.25mM EDTA to stop the reaction, and immediately loaded the entire reaction volume into a 1.5% agarose gel. We note that we additionally ran DNA from A4 and from the F_1_ progeny of A3 × A4 crosses (which will be heterozygous for the 22-bp InDel) through this genotyping pipeline as negative and positive controls, respectively.

Our CRISPR/Cas9 editing was reasonably successful and we generated 16 strains with homozygous third chromosomes carrying independent mutations in *Ugt86Dd* (File S4). Simultaneously, we generated seven strains containing un-edited homozygous third chromosomes. Our rationale for keeping such genotypes was that during the creation of our custom injection strain, and while establishing homozygous edited chromosomes, we had employed several generations of crossing with balancers. Since movement of genetic material via gene conversion from a balancer to the non-balancer homolog has been inferred (Blumenstiel *et al.* 2009), non-edited genotypes may prove useful controls since they have been through many of the same crosses as edited genotypes. Each of the 23 strains created was phenotyped using 5-6 replicate no-drug and nicotine-containing vials.

For two edited, and one un-edited genotype we used standard fly crosses to substitute the edited third chromosome into the A4 background, allowing a direct comparison of mutated and wildtype versions of *Ugt86Dd*. Each of these strains - A4-*Ugt86Dd*^*Del1*^, A4-*Ugt86Dd*^*Del11*^, and A4-*Ugt86Dd*^*wt*^ (File S4) - was tested directly as an inbred genotype, and as a heterozygote following crosses to A4, using 10 replicate no-drug and nicotine-containing vials.

### Estimating the frequency of the InDel in nature

Since the InDel is just 22-bp, we reasoned we could identify the insertion and deletion alleles directly from reads resulting from the next-generation sequencing (NGS) of pooled population samples of *D. melanogaster*, thereby estimating allele frequency. We retrieved raw FASTQ files from 13 wild-caught, resequenced population samples (Bergland *et al.* 2014) via the NCBI SRA (BioProject accession PRJNA256231), and used a simple UNIX grep procedure to count the number of reads containing the insertion and deletion sequences. We ensured that instances of a sequence occurring in both reads of a single paired-end fragment were counted only once. The insertion-specific sequence was taken as the central 20bp (i.e., ATTGTGGAGGACATTCATCG) of the 22-bp InDel, and its reverse complement. The deletion-specific sequence was taken as the 10bp either side of the 22-bp InDel (i.e., GAACGAATTCACTTCGTAGT), and its reverse complement. These sequences match all 15 inbred DSPR founder strains sequenced for the region (11 insertion lines and 4 deletion lines, Figure S1).

### Data availability

All raw phenotypic data associated with the study is presented in File S1. Raw RNAseq reads are available from the NCBI SRA (Accession SRP102254).

## RESULTS

### Large, dominant effect of chromosome 3 on nicotine resistance

Marriage et al. (2014) identified four autosomal QTL contributing to nicotine resistance in the pA DSPR population. Three QTL on chromosome two (Q1, Q2, Q3) individually contributed 4.6-8.5% to the broad-sense heritability of the trait, and one QTL on chromosome three (Q4) contributed 50.3% to the heritability. Based on estimating the effects associated with each founder allele at each mapped QTL it appeared that in each case carrying an A4 allele at the QTL led to greater resistance than carrying an A3 allele. To confirm this we tested various chromosome substitution genotypes, and observed that substituting the third chromosome from A4 into an otherwise A3 background leads to a marked increase in resistance compared to that exhibited by founder A3 (*P* < 10^−8^, Figure 1). This likely highlights the large effect of Q4, suggesting that A3 and A4 do harbor different alleles for the causative variant(s). Notably the effect of the A4 third chromosome appears to be dominant, with just one copy of the A4 chromosome in an A3 background sufficient to entirely recapitulate the phenotype of founder A4 (Figure 1).

**Figure 1.**
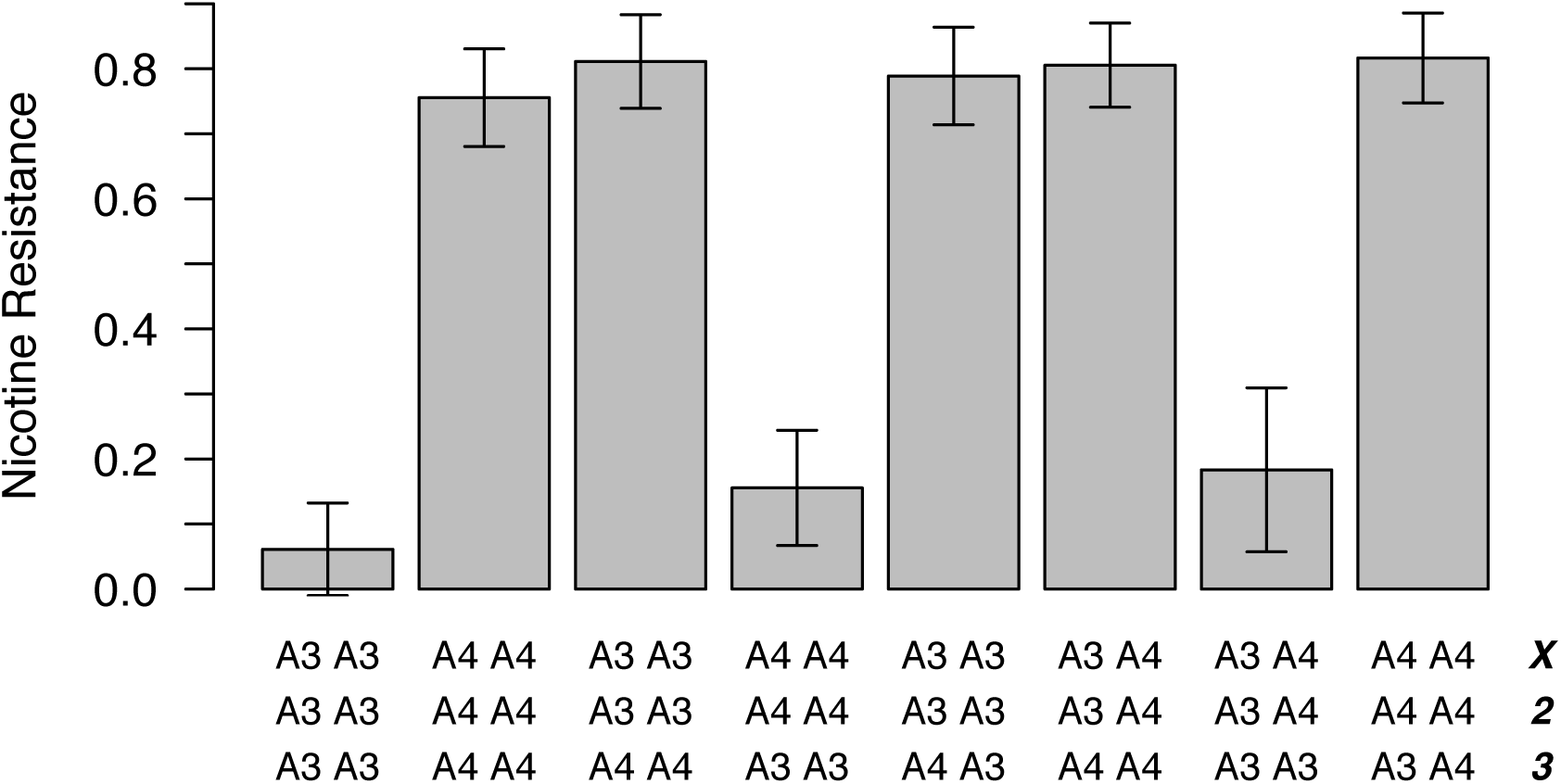
Effects of chromosome substitution on nicotine resistance. Starting from DSPR founders A3 and A4 we created a number of homo-and heterozygous chromosome substitution genotypes. The nicotine resistance phenotype of each genotype was tested across six replicate vials, and the plot shows the mean (± 1-SD) of each. We carried out all possible pairwise *t*-tests between genotypes; The three low resistance genotypes are statistically indistinguishable, as are the five high resistance genotypes, while the low and high resistance genotypes are significantly different (*P* < 0.0001 in all cases). Note that the viability of each genotype was tested twice under no-drug conditions, and all genotypes showed a mean viability > 0.9.

In contrast to the large effect of chromosome three, the combined effects of the X and chromosome two on nicotine resistance are relatively small; A genotype with the A3 third chromosome in an otherwise A4 background is only marginally more resistant than founder A3 (*P* = 0.07, Figure 1). This result confirms the smaller phenotype effects estimated at the second chromosome QTL by Marriage et al. (2014).

### Fine mapping nicotine resistance QTL

One strategy to fine map loci that have been more broadly mapped using a linkage mapping approach is to employ quantitative complementation tests, either using loss-of-function mutant alleles of plausible candidate genes (Long *et al.* 1996), or deletions that remove sections of the genome and tens to hundreds of genes (Pasyukova *et al.* 2000). Here, we crossed two DSPR founder inbred strains that differ markedly in nicotine resistance, A3 and A4, to a series of mutation-carrying strains and their corresponding co-isogenic, mutation-free control strains, generating four types of progeny (A3/mutation, A3/control, A4/mutation, and A4/control). We then used a statistical test that estimates the effects of founder and mutation, and the interaction between these factors. A significant interaction - a quantitative failure to complement - implies the effects of founder alleles are different in the mutant and control backgrounds, and thus suggests the founders segregate for functional variation at the tested gene/deficiency.

We employed two small deficiencies that deleted overlapping regions of the Q1 region, one of which eliminates both of the P450 genes within the QTL interval, *Cyp28d1* and *Cyp28d2*, and one of which deletes only *Cyp28d1*. Since the deletions remove a total of 11-13 genes, including three additional P450s not within the Q1 interval, we also employed a pair of Minos insertions (Metaxakis *et al.* 2005) that insert within coding exons of *Cyp28d1* and *Cyp28d2*, likely disrupting gene function. In all four cases there was a highly significant Founder × Mutant interaction (*P* < 10^−5^, Figure S2) with the difference between A3 and A4 being much greater in the mutant background. Collectively these results imply that both *Cyp28d1* and *Cyp28d2* show functional differences between A3 and A4 that confer effects on phenotype.

We carried out a similar experiment for the Q4 region, testing three overlapping deficiencies that individually delete 11-19 genes, including variable numbers of the UGT genes resident within the Q4 interval. All three deletions exhibit a significant quantitative failure to complement (*P* < 10^−6^, Figure S2). Since *Ugt86Dc* is the only gene deleted by all three deletions a parsimonious explanation is that A3 and A4 have distinct nicotine resistance alleles at this gene. However, Marriage et al. (2014) showed that expression of this gene is reduced in A4 compared to A3, which is not the pattern one might expect under the assumption that *Ugt86Dc* leads to the enhanced nicotine resistance of A4. Additionally, Marriage et al. (2014) provided evidence that the founder allelic effects at Q4 do not fall into two groups, as would be expected if a single causative gene was responsible for the QTL. Thus, the different deletions may be uncovering variants in independent genes that affect phenotype.

Finally, we tested a Minos element in a coding exon of *Ugt86Dj*, and one that resides within the 3’UTR of one of the two isoforms of *Ugt86Dh*. (Mutations in a known background were only available for 2/10 of the UGT genes under Q4 when this experiment was conducted.) While we observed a significant Founder × Mutant interaction for both genes (*Ugt86Dj*, *P* < 0.001; *Ugt86Dh*, *P* < 0.01; Figure S2), in both cases the difference between the A3 and A4 alleles was greatest in the mutant background. These results suggest the effects are more likely due to epistasis than allelic failure to complement (see Mackay 2001; Geiger-Thornsberry and Mackay 2004), and do not provide strong evidence for the role of functional variation at *Ugt86Dh* or *Ugt86Dj* in resistance to nicotine.

### Candidate nicotine resistance genes via RNAseq

Under the assumption that some fraction of complex trait variation is regulatory in origin (Gibson and Weir 2005; Gilad *et al.* 2008; Gusev *et al.* 2014; Torres *et al.* 2014; Albert and Kruglyak 2015), we might expect to see changes in the expression of loci harboring causative loci. Marriage et al. (2014) used RNAseq of whole, first-instar larvae from A3 and A4 tested under no-drug and nicotine-exposure conditions to attempt to resolve candidate genes. Here, we carried out a similar study employing mixed pools of RNA from relatively susceptible and relatively resistant pB DSPR RILs (File S2). An array of genes showed differential expression between susceptible and resistant genotypes and/or between treatments (File S5), but those changes at loci within QTL intervals are of principal interest.

Considering those 34 genes within Q1 only *Cyp28d1* was differentially-expressed at a nominal 5% level, showing an induction in expression on nicotine exposure in both susceptible (*P* < 10^−4^) and resistant animals (*P* = 0.001), confirming the results of Marriage et al. (2014). However, unlike our previous study we found no change in expression between susceptible and resistant pB line pools that would indicate an allelic difference between genotypes at this gene. Since Q1 was not identified in the pB DSPR panel it is possible there is no segregating variation that leads to expression variation within this panel, although we cannot discount the possibility we failed to capture any such variation in the small number of pB lines used for the current expression study (File S2).

At Q4, 5/10 UGT genes show a change in expression in at least one contrast, with two genes showing a change between susceptible and resistant genotypes. *Ugt35b* shows a slight increase in expression in the resistant genotypes under no-drug conditions (*P* < 0.05), while *Ugt86Dd* increases in expression in resistant animals under both no-drug and nicotine conditions (*P* < 10^−4^ and *P* = 0.001, respectively). Marriage et al. (2014) found no effect at *Ugt35b*, but did observe expression differences between A3 and A4 in the expression of *Ugt86Dd*. Since Q4 was identified in both the pA and pB populations, the similar change in expression between susceptible and resistant genotypes derived from the two panels implies *Ugt86Dd* is a strong candidate to harbor functional genetic variation contributing to resistance.

Notably, Marriage et al. (2014) identified a small peak in the LOD score profile in pB that encompasses the *Cyp6g1* gene, a gene shown to be associated with DDT (Daborn *et al.* 2002; Chung *et al.* 2007; Schmidt *et al.* 2010) and nicotine resistance (Li *et al.* 2012). However, we did not focus on the QTL in our previous study given the modest LOD score at the peak, and the relatively small number of pB RILs we assayed. Here, we found a dramatic increase in expression of *Cyp6g1* between susceptible and resistant pB genotypes in both no-drug and nicotine conditions (*P* < 10^−4^) providing some additional support for the effect of this gene on nicotine resistance in the DSPR.

### Significant reduction in resistance following ubiquitous gene knockdown

Under Q1 the genes most likely to harbor functional variation impacting nicotine resistance are *Cyp28d1* and *Cyp28d2*. Under Q4, of the ten UGT genes *Ugt86Dd* appears to be a strong candidate based on RNAseq data. We knocked down the expression of these three plausible candidates using 2-3 different UAS transgenes per gene. In comparison with co-isogenic control strains, we see a robust reduction in resistance following knockdown of each gene (Figure 2), suggesting these genes are functionally important for resistance. Notably, gene knockdown had no effect on viability under no-drug conditions; All strains showed phenotypes above 0.93 on no-drug food (compare to Figure 2), and there was no effect of gene knockdown under no-drug conditions (*P* > 0.4 for all tests). Thus, the effects of reducing expression of *Cyp28d1*, *Cyp28d2*, and *Ugt86Dd* appear to be specific to nicotine resistance.

**Figure 2.**
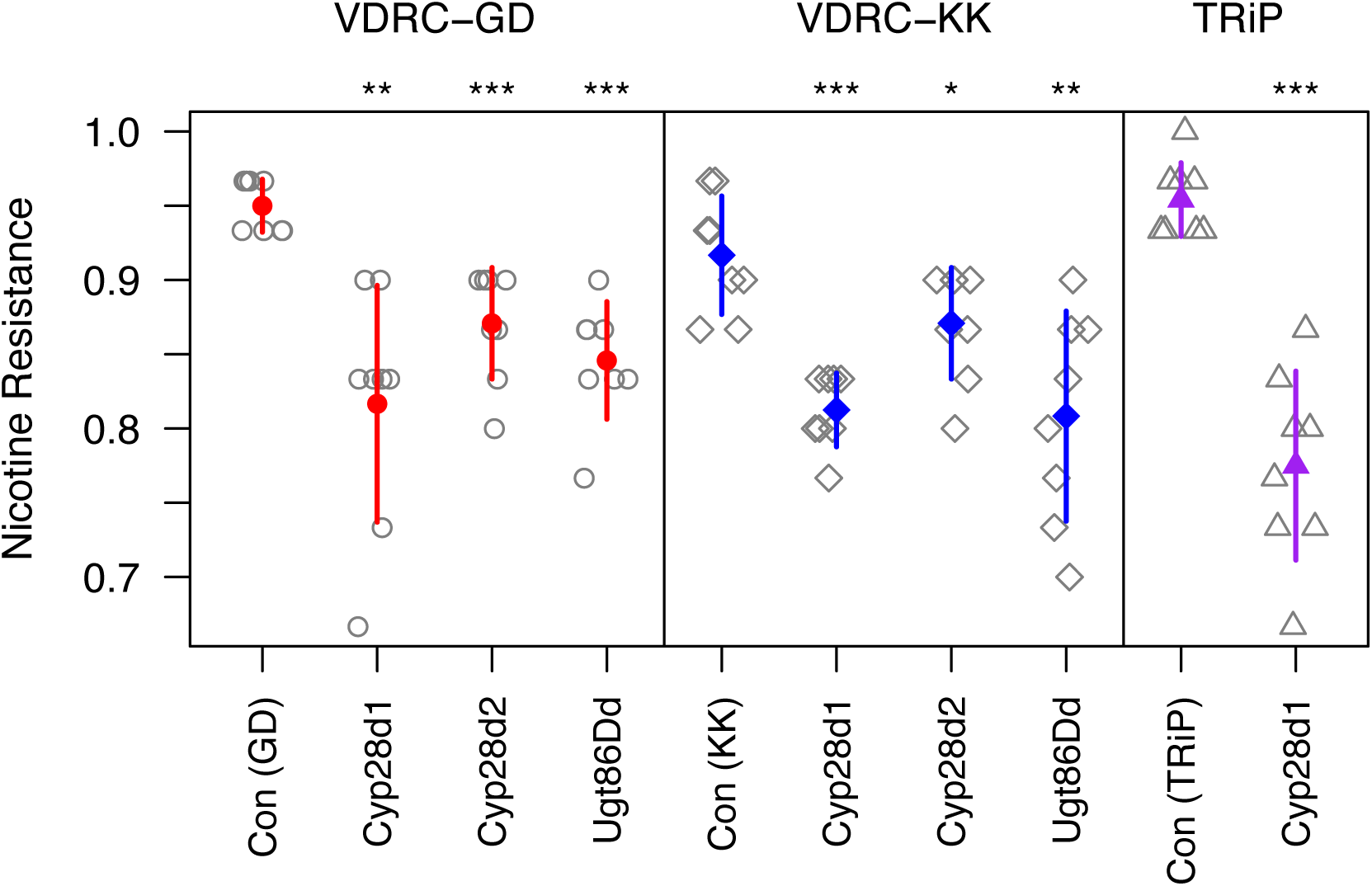
Effect of ubiquitous gene knockdown on nicotine resistance. We employed the Gal4-UAS-RNAi system to knock down the expression of three genes in all cells and at all timepoints via *Act5C*-Gal4. Each genotype was tested across eight replicate vials (raw data shown in gray symbols), and the mean (± 1-SD) phenotype of each is shown with colored symbols/lines. Within each RNAi system (VDRC-GD, VDRC-KK, TRiP) we tested the effect of each gene knockdown against its co-isogenic control using *t*-tests, and significance is highlighted by asterisks (* = *P* < 0.05, ** = *P* < 0.01, *** = *P* < 0.001). All ubiquitous RNAi knockdowns reduce nicotine resistance. Note that the viability of each genotype was tested 4-5 times in no-drug conditions, and all genotypes showed a mean viability of > 0.93.

### Knockdown of *Cyp28d1* and *Ugt86Dd* in the midgut reduces resistance

To help determine the tissue in which expression reduction leads to a decrease in nicotine resistance we used a series of tissue-specific Gal4 drivers. Using two drivers that express Gal4 in the anterior region of the midgut we saw that RNAi of *Cyp28d1* (in 5/6 cases) and *Ugt86Dd* (in 4/4 cases), but not *Cyp28d2* (in 0/4 cases), led to a significant reduction in resistance (Figure 3). In contrast, a driver expressing Gal4 in the posterior region of the midgut showed no effect of any target gene (Figure 3). These data imply that the anterior midgut is an important site of nicotine metabolism, and is impacted by the action of the products of *Cyp28d1* and *Ugt86Dd*, genes showing expression variation between resistant and susceptible genotypes (above and Marriage *et al.* 2014). The lack of any apparent effect of *Cyp28d2* in the midgut (Figure 3), in contrast to the effect observed following ubiquitous knockdown of the gene (Figure 2) might imply that *Cyp28d2* is required in a different, and un-tested tissue. Alternatively, the ubiquitous knockdown effect described above may simply be a false positive, which are known to occur at low rates in large-scale RNAi screens (e.g., Mummery-Widmer *et al.* 2009; Schnorrer *et al.* 2010).

**Figure 3.**
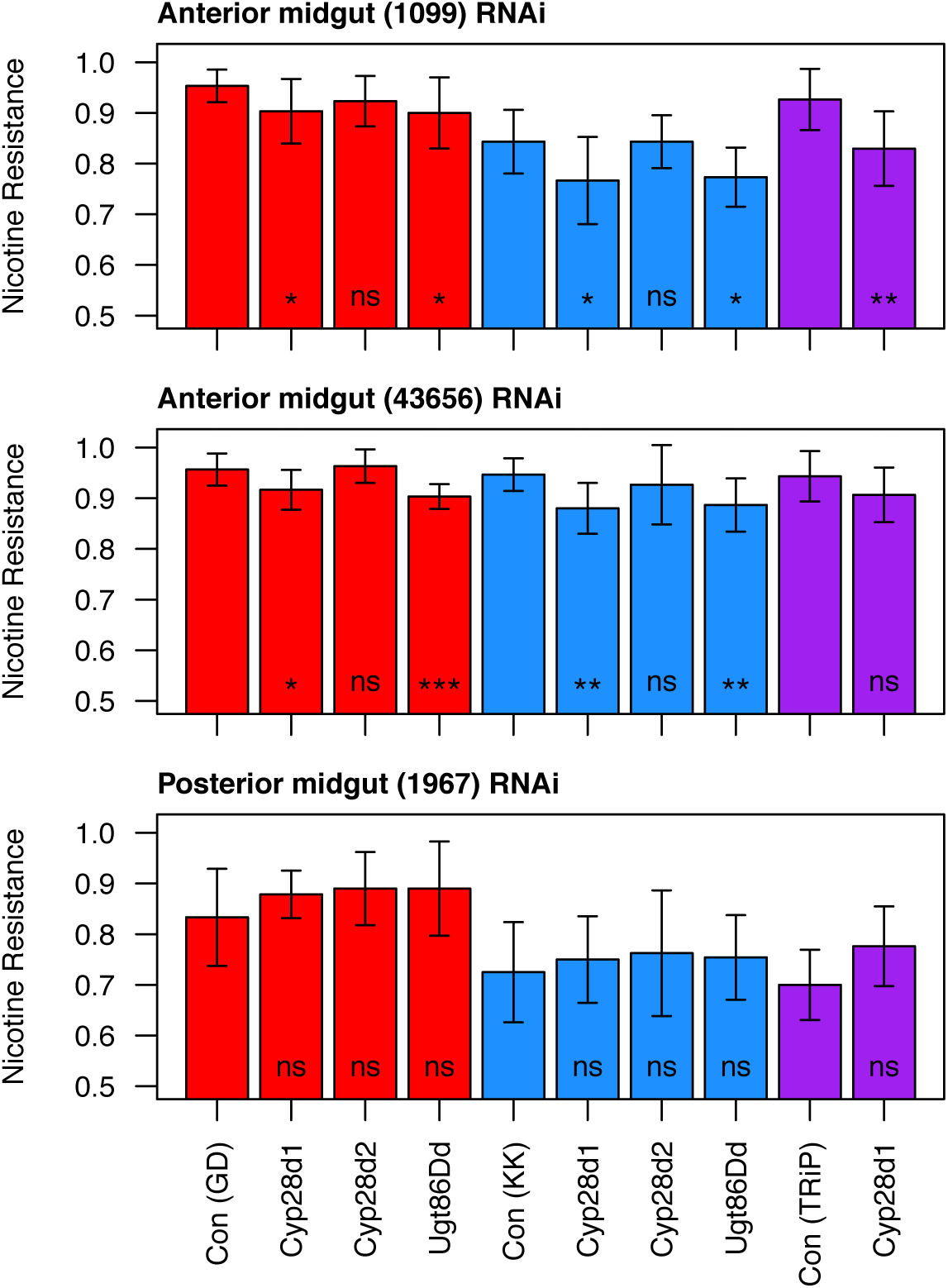
Effect of midgut-specific gene knockdown on nicotine resistance. We employed the Gal4-UAS-RNAi system to knock down expression of three genes in the midgut. Each genotype was tested across 7-10 (mean = 9.5) replicate vials, and we plot the mean (± 1-SD) for each. We used three RNAi systems (VDRC-GD = red, VDRC-KK = blue, TRiP = purple) and tested the effect of each gene knockdown against its co-isogenic control using *t*-tests; Significance is highlighted by asterisks (ns = P ≥ 0.05, * = *P* < 0.05, ** = *P* < 0.01, *** = *P* < 0.001). In all but one case, reduction of *Cyp28d1* and *Ugt86Dd* expression in the anterior midgut reduces resistance. The viability of each genotype in the plot was tested 2-5 times in no-drug conditions; These genotype means averaged 0.92, 24/30 of the genotypes have greater no-drug viability than nicotine viability, and for the other six genotypes no-drug viability was at most 4% less than that on nicotine food. These observations suggest the effects of these tissue-specific gene knockdowns are not a result of general defects in viability.

### Malpighian tubule knockdown of target detoxification genes enhances resistance

The malpighian tubules are an important site of xenobiotic metabolism in insects (Dow and Davies 2006; Yang *et al.* 2007), and we tested multiple Gal4 drivers that broadly express in the malpighian tubules, and also those that target specific cell types within the organ. RNAi against three genes (*Cyp28d1*, *Cyp28d2*, *Ugt86Dd*) using drivers expressing Gal4 broadly in the malpighian tubules, and in one case additionally in the hindgut and ureter, revealed no major changes in resistance (Figure 4). However, in nearly all cases knockdown of these genes in either principal or stellate cells of the malpighian tubule led to a significant *increase* in resistance in comparison with control genotypes (Figure 4). We note that under no-drug conditions all genotypes associated with these malpighian tubule cell-specific tests show viabilities above 0.93, implying the relative increase in resistance on gene knockdown is in response to the nicotine treatment.

**Figure 4.**
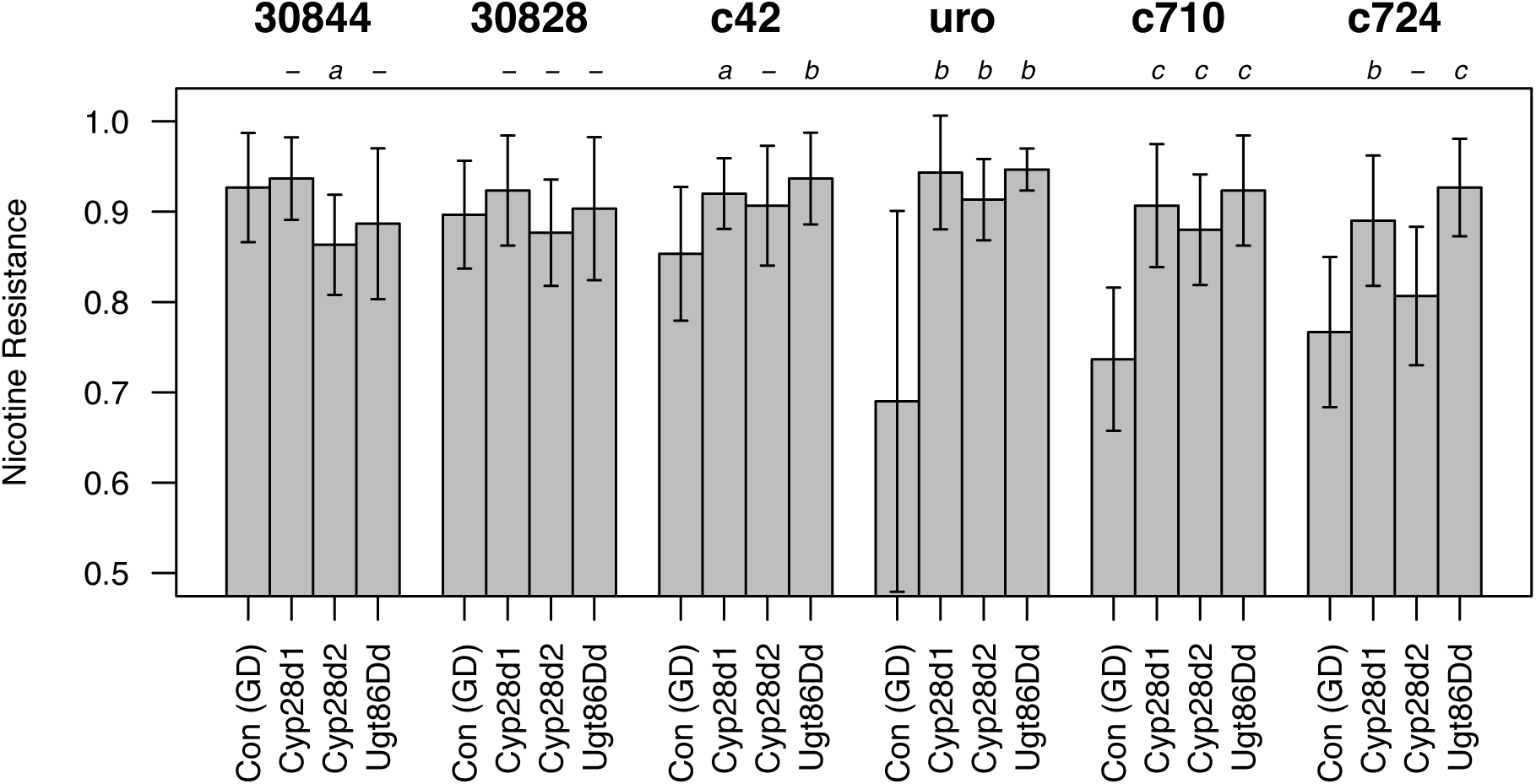
Effect of malpighian tubule RNAi on nicotine resistance. We knocked down expression of three genes (using UAS constructs from the VDRC-GD collection) with six different Gal4 drivers. The 30844 driver expresses Gal4 in the hindgut, ureter, and malpighian tubules, 30828 expresses Gal4 in the malpighian tubules, c42-Gal4 and uro-Gal4 are specific to the principal cells of the malpighian tubules, and c710-Gal4 and c724-Gal4 are specific to tubule stellate cells. Each genotype was tested across 10 replicate vials, and we plot the mean (± 1-SD) for each. The effect of each gene knockdown was compared to the relevant co-isogenic control using *t*-tests; Significance is highlighted by letters (-= *P* ≥ 0.05, *a* = *P* < 0.05, *b* = *P* < 0.01, *c* = *P* < 0.001). The viability of each genotype in the plot was also tested 2-5 times under no-drug conditions. The lowest genotype mean under no-drug conditions was 0.88, and the average genotype mean was 0.95. Notably, the mean phenotypes under no-drug conditions for the non-RNAi, control genotypes - marked “Con (GD)” in the plot - were 0.91-0.98, implying that the marked reduction in viability for some of these genotypes on nicotine-supplemented media is not due to general viability defects.

This result - heightened resistance to a drug after knocking down the expression of known detoxification genes in cell types known to be involved in xenobiotic metabolism - is counterintuitive. Indeed, RNAi knockdown of the P450 gene *Cyp6g1* in principal cells robustly leads to a reduction in DDT resistance (Yang *et al.* 2007). We speculate that the presence of nicotine or its byproducts, coupled with a reduction in expression of key nicotine metabolizers in the malpighian tubule principal/stellate cells, leads to the production of other detoxification enzymes that enhance resistance, but in a way that is mechanistically distinct from that observed in other genotypes. Analyses of gene expression specifically in the malpighian tubules of these knockdown genotypes may provide insight into this observation.

### Overexpression of functional *Ugt86Dd* enhances nicotine resistance

Using the *Ugt86Dd* gene sequence from the relatively susceptible A3 founder and the relatively resistant A4 founder, we made a series of UAS-*Ugt86Dd* strains allowing ectopic overexpression of the Ugt86Dd gene product. In the course of verifying clones prior to plasmid injection we noticed that exon 2 of the A3 allele contained a 22-bp deletion relative to A4, leading to an frameshift and a premature stop codon reducing the length of the predicted protein product from 517 amino acids to 206. PCR and sequencing of the original founders revealed that the A3 and A4 strains differ at this variant.

Ubiquitous overexpression of UAS-*Ugt86Dd*^*A4*^ via *Act5C*-Gal4 did not yield viable Gal4-UAS offspring, in contrast to similar overexpression of UAS-*Ugt86Dd*^*A3*^, implying both that overexpression of *Ugt86Dd* is poisonous to cells, and that A3 (a strain that is relatively susceptible to nicotine) may carry a nonfunctional allele of *Ugt86Dd*. To test the effect of *Ugt86Dd* overexpression we used a series of drivers expressing Gal4 in various regions of the gut, and compared a single line containing the UAS-*Ugt86Dd*^*A3*^ transgene to five strains containing the same UAS-*Ugt86Dd*^*A4*^ transgene. Figure 5 shows that for every Gal4 driver, overexpression of UAS-*Ugt86Dd*^*A4*^ leads to higher nicotine resistance than overexpression of UAS-*Ugt86Dd*^*A3*^. These data suggest that *Ugt86Dd* is an excellent candidate to contribute to nicotine resistance, and further suggests that the InDel variant segregating between strains A3 and A4 may be responsible for some of the difference in phenotype exhibited by these strains. We note that while all six strains tested are homozygous for the same third chromosome harboring the transgene landing site, they may be variable on the second and X chromosomes due to post-injection crossing against balancers. So it is conceivable that some of the variation observed between the A3 and A4 transgenes is due to differences in genetic background.

**Figure 5.**
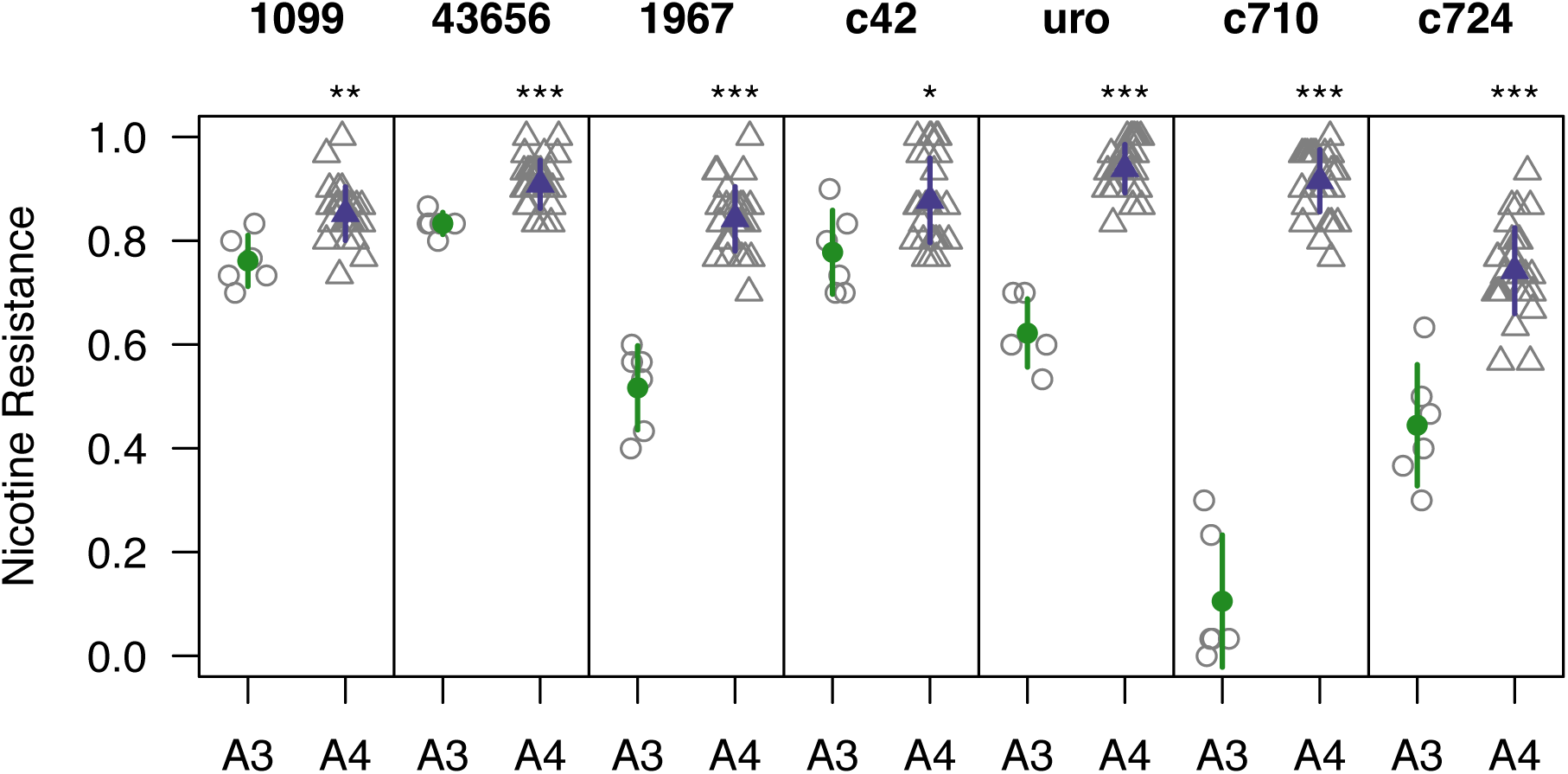
Effect of tissue-specific overexpression of *Ugt86Dd* alleles. We overexpressed *Ugt86Dd* derived from strains A3 (which contains a 22-bp out-of-frame coding deletion) and A4 (which carries an allele making full length gene product) using seven tissue-specific Gal4 drivers. We had access to a single UAS-*Ugt86Dd*^*A3*^ transgene line and five UAS-*Ugt86Dd*^*A4*^ transgene lines, and tested each line over six replicate vials (gray open symbols). We present the mean (± 1-SD) phenotype of each genotype in colored symbols, and compared the genotype means for each Gal4 driver using *t*-tests (* = *P* < 0.05, ** = *P* < 0.01, *** = *P* < 0.001). In every case the A4 transgene leads to higher resistance. Notably, we additionally tested all genotypes across 2-3 no-drug vials, and the A3 transgene showed viabilities 0.84-0.96, indicating that the marked reductions in viability under nicotine conditions is nicotine-specific.

This overexpression experiment may not say anything about the natural site of *Ugt86Dd*-based nicotine detoxification, since we are ectopically expressing the gene in tissues where it may not normally be expressed under its native promoter.

### The *Ugt86Dd* InDel variant is associated with nicotine resistance

We used PCR and Sanger sequencing to show that four of the DSPR founder strains (A3, AB8, B6, and B7) possess the 22-bp deletion allele at *Ugt86Dd* (Figure S1). At the Q4 QTL identified by Marriage et al. (2014) these four founders have the lowest strain effects, suggesting the InDel may have a functional role in resistance. Furthermore, mean phenotypes of RILs carrying the insertion and deletion alleles are highly significantly different for both the pA and pB populations (*t*-test, *P* < 10^−15^; File S3), with the mean resistance of the deletion-containing RILs being 37% (pA) and 27% (pB) less than that of the insertion-containing RILs.

We employed InDel status as an additional covariate in the DSPR QTL mapping analysis and succeeded in substantially reducing the LOD score at the site of the Q4 locus (Figure 6), eliminating any above-threshold peak in pB, and leaving a much more modest effect at the locus in pA; In the standard analysis Q4 contributes 46.5% to the variation among pA lines, whereas the above-threshold peak at the same location after accounting for the InDel contributes 5.3% to the variation. These data imply that the InDel variant alone, or one or a collection of variants in linkage disequilibrium (LD) with this variant are responsible for a large fraction of the nicotine resistance variation in the DSPR, and may completely, or nearly completely explain the major-effect QTL mapped by Marriage et al. (2014).

**Figure 6.**
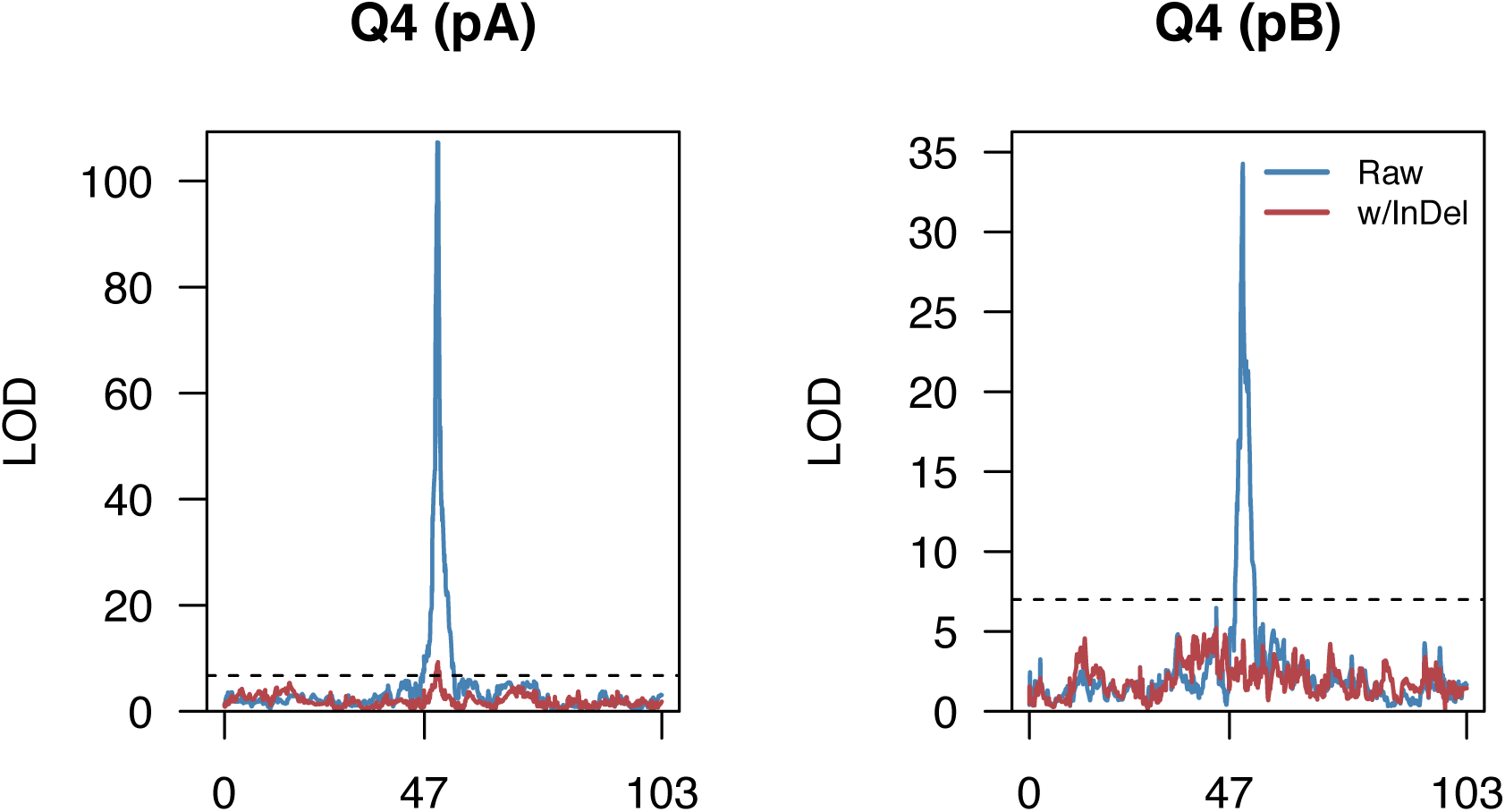
An InDel variant in *Ugt86Dd* explains variation at a nicotine resistance QTL. Each plot shows the LOD curve along chromosome three, with genetic distance along the *x*-axis, in the pA (left) and pB (right) DSPR populations. In blue, using a standard mapping analysis in the DSPR, one sees evidence for the major-effect QTL Q4. In red, using a similar analysis but now including a covariate accounting for the InDel status of each phenotyped RIL, one sees a peak in pA with a LOD score just surviving the threshold, and no peak in pB. These data suggests the InDel variant may contribute to the effect observed at Q4.

To attempt to validate this association, and specifically test the effect of the InDel variant on nicotine resistance, we exploited the DGRP, a collection of around 200 inbred, re-sequenced strains of *D. melanogaster* (Mackay *et al.* 2012; Huang *et al.* 2014). Seven of the DGRP strains were found to harbor the same 22-bp deletion allele as found in the DSPR. We intercrossed these lines for seven generations to generate a population fixed for the deletion allele, but otherwise outbred, and generated a similar population from a random set of seven DGRP strains carrying the insertion allele. Testing these two populations revealed a small, yet significant reduction in resistance in the deletion population (mean of insertion population = 0.94, mean of deletion population = 0.89, *t*-test *P* = 0.011). While this result does tend to validate our work in the DSPR, any effect of the *Ugt86Dd* InDel variant is apparently much smaller in the DGRP.

### CRISPR/Cas9-derived deletion mutations at *Ugt86Dd* in A4 markedly reduce resistance

The effect of the deletion allele in founder A3 is to generate a premature stop codon in *Ugt86Dd*. To directly test the effect of premature stop-encoding mutations in this gene, we generated a series of deletion-alleles via CRISPR/Cas9 editing. Using standard fly crosses with balancers we constructed a custom injection strain containing an X-linked *vasa*-Cas9 transgene and an A4-derived third chromosome, and employed a guide RNA that generates double-strand breaks at the site of InDel (Figure S1). We generated a series of 16 independent mutations (File S4), 13 of which lead to premature stop codons, and constructed strains carrying homozygous edited third chromosomes for all 16. Simultaneously, we made seven strains with homozygous un-edited third chromosomes from genotypes that were passed through the CRISPR/Cas9 process (injection, balancing, and so on). Since it is known that the sequence of the non-balancer homolog can be altered by gene conversion from the balancer (Blumenstiel *et al.* 2009), such un-edited genotypes provided valuable controls for the effects of the induced-mutations. All 23 strains were tested in multiple replicates for viability under no-drug and nicotine conditions, and while we observed no difference between the sets of un-edited and edited genotypes in the absence of nicotine (*P* = 0.2), we saw a marked decrease in resistance due to editing (*P* < 10^−15^, Table S1).

The fact that the three lesions that result in amino acid changes, but do not introduce a premature stop codon, still each lead to a strong reduction in nicotine resistance may suggest that the region of the protein targeted for mutation is critical to the function of Ugt86Dd.

For two deletion alleles - a 1-bp deletion and an 11-bp deletion - that both lead to premature stop codons, along with one un-edited allele, we used standard fly genetics to put the third chromosome (originally derived from A4) into a complete A4 background, generating strains A4-*Ugt86Dd*^*Del1*^, A4-*Ugt86Dd*^*Del11*^, and A4-*Ugt86Dd*^*wt*^. All three have marginally reduced viability under no drug conditions compared to the original A4 founder strain (*P* = 0.03-0.06), perhaps indicative of novel mutations relative the A4 progenitor line, or slight changes in sequence due to gene conversion from balancers during stock construction. Nonetheless, Figure 7 shows that both of the deletion-carrying stocks A4-*Ugt86Dd*^*Del1*^ and A4-*Ugt86Dd*^*Del11*^ have significantly reduced nicotine resistance in comparison with A4-*Ugt86Dd*^*wt*^ (*P* < 10^−4^ and *P* < 0.001, respectively). The CRISPR/Cas9-induced mutations appear to behave as recessive alleles since measuring the phenotype of their heterozygous progeny after crossing to A4 recapitulates the A4 homozygous phenotype (Figure 7).

**Figure 7.**
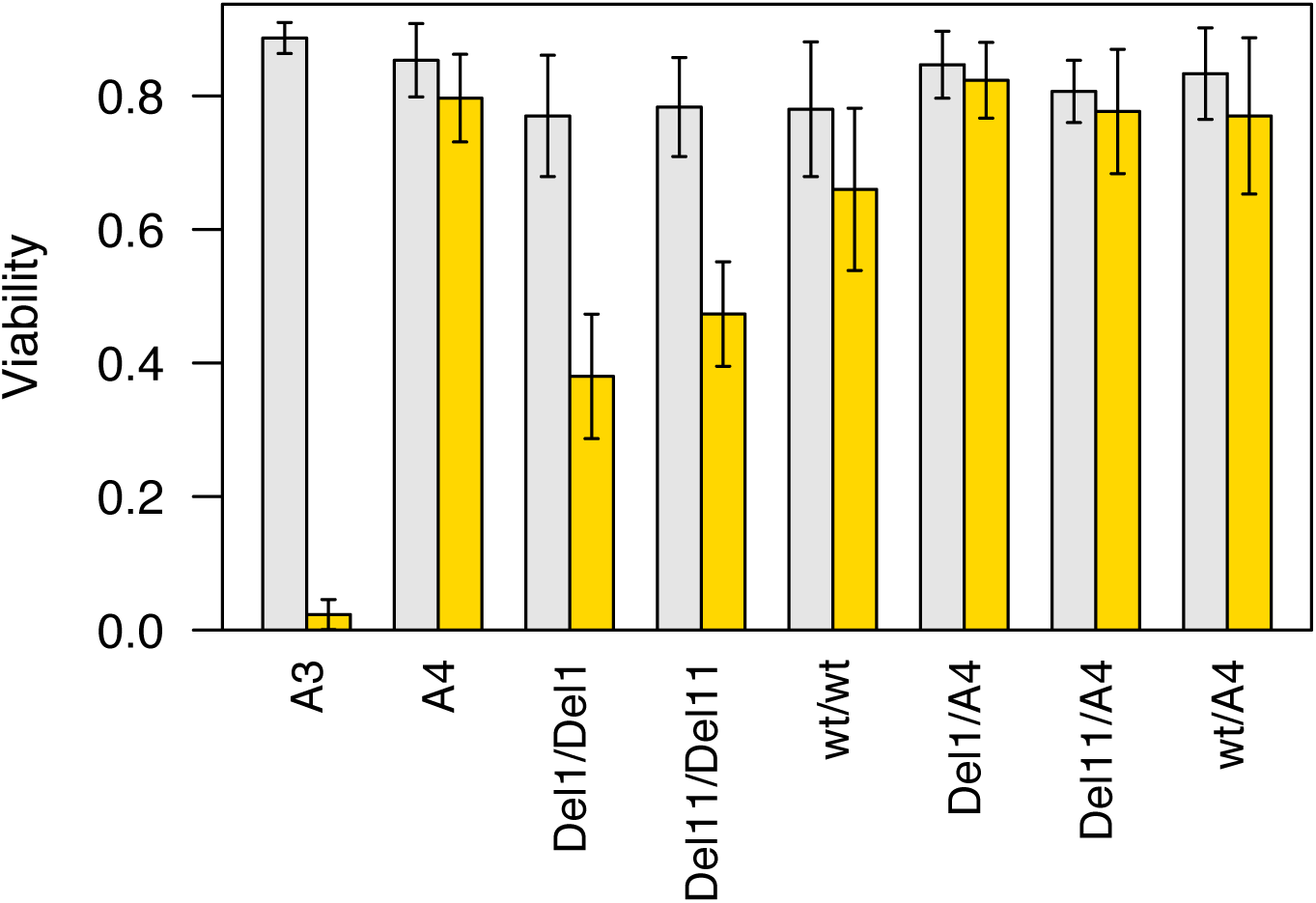
Phenotypes of background-controlled *Ugt86Dd* CRISPR/Cas9-derived mutations. We made a pair of *Ugt86Dd* mutations homozygous in the A4 background. *Del1* introduces a 1-bp mutation, while *Del11* introduces an 11-bp mutation, each leading to a different stop codon mutation (File S4). In addition, we passed an un-edited allele (*wt*) through an identical crossing scheme, which should be identical to the pure A4 strain. We tested 10 replicate vials of each of the genotypes in the plot for both no-drug (gray bars) and nicotine (yellow bars) treatments, and present the mean (± 1-SD) viability across replicate vials. The two homozygous mutant strains (*Del1*/*Del1* and *Del11*/*Del11*) have significantly lower nicotine resistance (*P* < 10^−5^) in comparison to the parental A4 genotype, and the un-edited control strain (*wt*/*wt*). Since the un-edited control strain has slightly, but significantly reduced resistance in comparison to A4 (*P* < 0.01), the crossing scheme used to establish the homozygous CRISPR/Cas9-derived genotypes may have resulted in some additional changes in addition to the target mutation.

The difference in nicotine resistance between A3 and A4 is 0.69-0.77 (Figures 1 and 7), while A4-*Ugt86Dd*^*Del1*^ and A4-*Ugt86Dd*^*Del11*^ differ from A4-*Ugt86Dd*^*wt*^ by 0.28 and 0.19, respectively. This suggests that 25-41% of the difference between founders A3 and A4 could be due to functional variation at *Ugt86Dd*, with the most likely variant conferring this effect being the naturally-segregating 22-bp InDel. Since the difference between the mean phenotype of RILs carrying the A3 and A4 alleles at the Q4 locus is 0.37 (Marriage *et al.* 2014), our gene editing data again implies that *Ugt86Dd*, and likely the InDel variant that results in an allele containing a similar premature stop codon to the edited mutations, confers the bulk of the effect observed at this QTL.

## DISCUSSION

### Variation at *Cyp28d1* may underlie nicotine resistance

Evidence from deficiency and insertional mutant quantitative complementation tests, coupled with prior RNAseq data (Marriage *et al.* 2014), and both ubiquitous and tissue-specific RNAi strongly implicates variation at *Cyp28d1* in the genetic control of nicotine resistance. Data from Chakraborty et al. (2017), who have generated an assembly of DSPR founder A4 based on long-read sequencing data, give some indication that variation in *Cyp28d1* copy number may be causative; In contrast to the *D. melanogaster* reference strain, founder A4 - which is relatively resistant to nicotine - possesses two copies of *Cyp28d1*. Marriage et al. (2014) showed that *Cyp28d1* expression is relatively high in A4, supporting the idea of a positive correlation between copy number and gene expression, which could in turn lead to enhanced resistance. Such positive associations between P450 copy number and xenobiotic resistance have been reported previously (Wondji *et al.* 2009; Schmidt *et al.* 2010). Nonetheless, the relationship between copy number and expression level is not straightforward (Zhou *et al.* 2011; Schrider *et al.* 2016), and while the presence of two copies of *Cyp28d1* in a strain resistant to nicotine is suggestive, establishing a direct effect clearly requires functional validation.

Evidence that *Cyp28d2* also contributes to nicotine resistance, and segregates for causative variation generating the Q1 locus, is marginally more mixed. A quantitative complementation test using an insertional mutant implies the gene harbors a functional difference between strains A3 and A4, strain A4 exhibits higher expression than A3 (Marriage *et al.* 2014), and ubiquitous knockdown of *Cyp28d2* decreases resistance. However, midgut-specific knockdown of the gene had no detectable effect in the present study. It is of course possible that we did not knock down *Cyp28d2* in the tissue in which it would normally act on nicotine, so failing to identify an effect. Employing an expanded catalog of Gal4 drivers (Buchon *et al.* 2013) may uncover the tissue in which *Cyp28d2* is active against nicotine. Given our data, either *Cyp28d1* or *Cyp28d2*, or indeed both genes, may contribute to the effect observed at the Q1 locus originally mapped by Marriage et al. (Marriage *et al.* 2014). Additional fine mapping in natural populations, where both genes appear to segregate for copy number variants (Good *et al.* 2014), may be profitable, and allow functional effects associated with the two genes to be separated.

### *Ugt86Dd* explains a large fraction of the variation for nicotine resistance in the DSPR

All of the evidence we present here and in Marriage et al. (2014), including quantitative complementation tests, RNAi, overexpression and CRISPR/Cas9 editing, strongly implicates *Ugt86Dd* as a major source of genetic variation contributing to nicotine resistance. The segregating 22-bp coding InDel is a compelling candidate for the actual causative site underlying much of this variation; By accounting for the polymorphism in the QTL mapping the effect of the Q4 interval is markedly reduced, and generating edited mutations with similar premature stop codons to that conferred by the naturally-occurring deletion results in a dramatic reduction in nicotine resistance.

Several pieces of evidence suggest that other polymorphisms additionally contribute to the effect observed at the Q4 locus. First, the strain effects at the QTL do not fall into two categories, as would be expected if a biallelic causative polymorphism was solely responsible for the QTL effect (Marriage *et al.* 2014). Second, a quantitative complementation test using a deficiency that eliminates several UGT genes, but leaves *Ugt86Dd* intact, showed a significant failure to complement, implying other genes in the QTL region may segregate for functional variation (Figure S2). Third, after accounting for the InDel, QTL mapping in the pA population still yields a modest-effect QTL at the Q4 interval. Finally, the difference between the strain effects of A4 (carries the *Ugt86Dd* insertion) and A3 (carries the deletion) at QTL Q4 is larger than the phenotypic difference between wildtype and mutant *Ugt86Dd* alleles in the same A4 background. One factor that could contribute to the Q4 QTL in addition to *Ugt86Dd* is a copy number variant at *Ugt86Dh* (Chakraborty *et al.* 2017), where founder A4 harbors two copies of this gene relative to the *D. melanogaster* reference genome (although we do not yet know the status of this gene in other DSPR founders.) Notably two copies of *Ugt86Dh* increases expression (Chakraborty *et al.* 2017), implying the CNV may have some functional effect, and the gene is deleted in the one deficiency that does not eliminate *Ugt86Dd*. Given the circumstantial evidence associated with *Ugt86Dh* it is a strong candidate for additional followup, functional characterization.

### A segregating InDel at *Ugt86Dd* likely contributes to nicotine resistance

Our QTL mapping re-analysis and overexpression studies implicate the naturally-segregating 22-bp InDel polymorphism in *Ugt86Dd* as a major causative factor in nicotine resistance in the DSPR. While we don’t specifically generate the 22-bp deletion in the A4 background, our creation and testing of other deletions at a similar location in the *Ugt86Dd* gene, that similarly lead to premature stop codons, provides strong evidence that the InDel is functional, and directly leads to an effect on phenotype.

Given the apparent large effect of the InDel in the DSPR, and specifically the effect of introducing a premature stop-encoding lesion into A4, we anticipated the effect of the deletion would be routinely large. However, comparison of DGRP-derived populations fixed for alternate alleles at the InDel, but otherwise outbred, showed only a modest effect of the variant. Our design assumes we can estimate the effect of each allele at the InDel averaged over a similarly randomized genetic background. However, due to the paucity of DGRP lines possessing the deletion, it is likely that the insertion and deletion populations show many other fixed differences along the genome, some of which could lead to underestimating any true effect of the variant.

Outside of this technical challenge, one additional possibility for the very different effect estimates of the InDel in DSPR-and DGRP-derived populations is a genetic interaction between allelic variation at the InDel, and that at other functional sites across the genome. Clearly such an assertion is speculative given the data we have available, but we know that epistasis contributes significantly to phenotypic variation (Monnahan and Kelly 2015; Forsberg *et al.* 2017), and large-effect mutations moved into a varied set of genetic backgrounds are known to result in a spectrum of phenotypic expression (Chandler *et al.* 2013; Chari and Dworkin 2013; He *et al.* 2014; Sittig *et al.* 2016). Thus, there is the potential for genetic background to influence the expression of the *Ugt86Dd* variant.

A way to both establish the causative effect of the 22-bp *Ugt86Dd* InDel in the DSPR, as well as to estimate any background-specific expression of the variant alleles in the DGRP (and other populations), is to use CRISPR/Cas9 to precisely introduce the deletion into strains carrying a wildtype *Ugt86Dd* sequence, and similarly “repair” the allele in strains carrying the deletion (see Gratz *et al.* 2013). Such reagents would additionally facilitate future work on the mechanisms underlying nicotine detoxification by *Ugt86Dd*, for instance by examining the products of nicotine metabolism (Hoi et al. 2014), and allow tests of the effect of the *Ugt86Dd* gene in the detoxification of other xenobiotics, perhaps neonicotinoid insecticides.

### *Ugt86Dd* deletion is rare and recessive

Among the 15 DSPR founder strains, a world-wide collection of *D. melanogaster* genotypes (King *et al.* 2012b), the 22-bp deletion allele at *Ugt86Dd* is fairly common (4/15 or 27%). It is less frequent in the DGRP, a series of inbred genotypes extracted from a single North American population (Mackay *et al.* 2012). In order to assess the frequency of the allele in multiple, wild-caught population samples, we searched for next-generation sequencing reads that perfectly match the deletion and insertion alleles at the variant in a series of pooled re-sequencing datasets derived from a number of US populations (see Materials and Methods and Bergland *et al.* 2014). Averaging over these samples we roughly estimate the frequency of the deletion in nature at around 2% (Table S2). There is fairly large sample-to-sample variation around this value (0-11%) that could represent real biological, among-population variation, or simply be a result of sampling (of both individuals and sequencing reads.)

The deletion allele leads to a premature stop, likely ablating gene function, and purifying selection may perhaps explain the low frequency of the allele in nature. Huang et al. (2014) identified variants in the set of 200 DGRP fly strains that potentially generate a damaged protein, and found that such variants are typically less frequent than other sites (e.g., non-synonymous variants). Similarly, resequencing of protein-coding genes in humans has shown that rare polymorphisms are enriched for putatively damaging events, such as nonsense mutations and frameshifts (MacArthur *et al.* 2012; Nelson *et al.* 2012; Tennessen *et al.* 2012). Since the *Ugt86Dd* deletion appears to be recessive (Figure 7), under the assumption there is some cost to an organism in nature carrying the deletion in homozygous form, this might explain why its frequency in nature is not even lower.

It is not clear why the deletion allele is fairly common in the DSPR founders. The founders were collected in the 1950’s and 1960’s from a number of different countries (King *et al.* 2012b), prior to the *P*-element sweep through *D. melanogaster*, and conceivably populations were enriched for the deletion during this period. However, given the small number of founders, and their arbitrary sampling from stock centers many generations after their original derivation from nature (King *et al.* 2012b), a deletion frequency of 4/15 may not reflect the actual worldwide population frequency of the allele at the time of collection. It is tempting to speculate that the frequency of the *Ugt86Dd* deletion allele has been modulated by the use of nicotine as an insecticide. Nicotine was in common use as a pesticide after World War II (Shepard 1951; Metcalf 1955), but is no longer used in this way in the US (Environmental Protection Agency 2009), having being supplanted by other insecticides, many of which are nicotine derivatives (Goulson 2013). However, it is known that individual UGT genes can act on a range of substrates (Tukey and Strassburg 2000), so any selection acting on *Ugt86Dd* could be due to one or more unknown compounds to which flies can be exposed in nature.

Despite the large effect of the InDel in the DSPR, replicated by our CRISPR/Cas9 mutations, the recessive nature of the deletion allele and its low frequency in nature likely means the InDel explains only a very small fraction of the variation in nicotine resistance in a natural, outbred diploid population. Using the formula appropriate for a completely recessive variant, *V*_a_ = 8*pq*^3^*α*^2^ (Falconer and Mackay 1996), estimating the frequency (*q*) of the deletion allele at 2%, taking the effect of the deletion from the difference between the background matched wildtype *Ugt86Dd* genotype and the CRISPR-derived lesions (0.19-0.28), and assuming the phenotypic variance among DSPR line means (Marriage *et al.* 2014) is representative of that in a natural population, the InDel explains less than 0.01% of the phenotypic variation for the trait. This estimate is lower still if the difference between homozygous insertion-and deletion-carrying genotypes are more in line with the values observed for the DGRP populations.

### Resolving causative polymorphisms underlying QTL

Several features of our study system facilitated the resolution of a likely causative sequence variant underlying a mapped QTL. First, our study of a xenobiotic resistance trait clearly motivated a focus on the small number of detoxification family members within QTL. This ability to hone in on genes that were *a priori* highly likely to harbor segregating causative variation may be unavailable for many other traits, where indeed the purpose of unbiased, genomewide mapping is to help elucidate the pathways underlying phenotypic variation. Second, we were fortunate that the causative gene possessed a variant with a clear molecular signature of damage. The phenotypic effects of functional nonsynonymous polymorphisms, and particularly those associated with noncoding changes, are considerably more challenging to decipher from sequence data alone. The relative ease with which certain types of coding change can be recognized explains in part why current catalogs of causative variants are biased towards sites in coding regions (Stern and Orgogozo 2008). Third, despite the QTL containing multiple detoxification genes, and our prediction that the QTL was generated by the action of multiple genes/alleles (Marriage *et al.* 2014), alone the *Ugt86Dd* coding mutation has a major effect on phenotype. This was unanticipated given previous work showing that broadly-mapped QTL fragment into multiple smaller-effect loci on fine mapping, but obviously facilitated detection of the polymorphism. Our result appears representative of the phenomenon that causative coding variants often have more severe effects on phenotype than do regulatory events, an observation most clearly seen in human Mendelian diseases where an overwhelming fraction of disease-causing lesions target coding DNA (Botstein and Risch 2003).

Tracking down the molecular lesion(s) underlying QTL contributing to trait variation is challenging, even in the *Drosophila* model system. In addition to Marriage et al. (2014), various studies have identified very large effect QTL with the DSPR (King *et al.* 2012b; Cogni *et al.* 2016), and for such QTL finding and validating the causative locus using the battery of tools available for *D. melanogaster* is within reach. However, QTL effects are typically more modest (Kislukhin *et al.* 2013; Highfill *et al.* 2016), often do not implicate unambiguous candidates, and may regularly be generated by noncoding variants (Albert and Kruglyak 2015). Such features ensure that some fraction of causative variants will always remain uncharacterized. Nonetheless, the increasing availability of powerful genomic and functional molecular genetics tools will allow many causative variants underlying important biomedical traits to be resolved. Building genome assemblies using long-read, single-molecule sequencing will help to generate a more complete catalog of variation segregating in mapping panels such as the DSPR (Chakraborty *et al.* 2017), potentially yielding strong candidate causative variants for experimental validation. Future experiments will additionally be able to take advantage of tissue specific RNAseq (e.g., King *et al.* 2014) and ATACseq (Buenrostro *et al.* 2013) in order to resolve putatively causative genes and regulatory regions, and help fine map functional noncoding variation (Kumasaka *et al.* 2016). Finally, the strengths of CRISPR/Cas9 genome editing to the eukaryotic quantitative genetics community cannot be overstated. For the first time we will be able to validate the effects of specific allelic variants in otherwise standardized genetic backgrounds, allowing subtle genetic effects to be accurately measured, and the downstream molecular effects of allelic substitutions to be precisely characterized *in vivo*.

## ACKNOWLEDGEMENTS

We thank Tony Long for suggesting the *in silico* InDel frequency estimation approach, Emily Pfeifer for technical assistance, and Rob Unckless for comments on a previous version of this manuscript. We thank the TRiP (NIH R01-GM084947) and the Vienna *Drosophila* Resource Center for providing transgenic RNAi fly stocks used in this study. Various additional fly stocks, including the DGRP strains, were obtained from the Bloomington *Drosophila* Stock Center (NIH P40-OD018537). We also thank Julian Dow and Shireen Davies (University of Glasgow) for providing malpighian tubule Gal4 driver strains, and Nicolas Buchon (Cornell) for donating a tissue-specific gut Gal4 driver (flygut.epfl.ch). Finally, we thank Johannes Bischof for donating the pUASg.attB vector used for the overexpression experiment (flyorf.ch). Xiaofei Wang’s work on this project was supported via the Kansas INBRE project (NIH P20-GM103418), and the KU Genome Sequencing Core is supported by an NIH COBRE (P20-GM103638). This project was supported by National Institutes of Health grant R01-OD010974 to S.J.M. and Anthony D. Long (UC Irvine).

## SUPPLEMENTAL DATA FILE LEGENDS

**File S1** Phenotype of every replicate vial scored in the study. A nine-column tab-delimited text file. The “Expt” column associates each replicate with the chromosome substitution study (‘ChrSub’), the quantitative complementation test study (‘QCT’), the ubiquitous and tissue-specific RNAi knockdowns (‘RNAi_ubiq’ and ‘RNAi_tissue’), the *Ugt86Dd* overexpression (‘OE’), the DGRP tests (‘DGRP’), and the CRISPR/Cas9 studies (‘CRISPR1’ examined all strains generated, and ‘CRISPR2’ examined only those mutations substituted into the A4 background). The “matGeno” and “patGeno” columns give the maternal and paternal genotypes of the test individuals (see Materials and Methods for strain IDs). The “Medium” column is either ‘nic’ (nicotine-containing food) or ‘con’ (food does not contain nicotine). The “Batch” and “RepVial” columns define the experimental batch the replicate vial was tested in, and assigns a number to each replicate vial. “NumLarvae” defines the number of first-instar larvae placed on the food, which in most cases is equal to 30, while “NumAdults” defines the number of adults that emerge from the vial. “Pheno” is the fraction of larvae that emerge as adults from each vial, and is the phenotypic score used throughout the study.

**File S2** The set of pB2 DSPR RILs used for RNAseq. RIL IDs are given as the standard DSPR nomenclature (see FlyRILs.org). The nicotine resistance phenotype data is taken directly from that presented in Marriage et al. (2012). The allele present in each RIL at the *Ugt86Dd* 22-bp InDel (In = insertion, Del = 22-bp deletion) is based on direct genotyping of the DSPR founder strains, and high confidence estimates of the founder contribution at the *Ugt86Dd* gene region in each RIL.

**File S3** The nicotine resistance phenotypes for all DSPR RILs used for re-mapping. A five-column tab-delimited text file. The “matRIL” and “patRIL” columns give the RIL IDs (see FlyRILs.org), “Pheno” is the mean nicotine resistance of the RIL, “SubPop” is the subpopulation of each RIL (1 = pA1 or pB1, 2 = pA2 or pB2; the first digit of each RIL ID gives the population of origin - 1 = pA, 2 = pB), and “InDel” states whether the RIL contains the insertion or deletion allele at the 22-bp *Ugt86Dd* InDel (1 = insert, 0 = deletion). Note that the data in this file is identical to that presented by Marriage et al. (2012) except that it only includes those 790 pA and 426 pB RILs for which we are confident of the InDel status.

**File S4** Sequences of all CRISPR/Cas9-generated mutations in *Ugt86Dd*. For each of the 16 mutations we give the DNA sequence, and the anticipated resulting amino acid sequence. The genotype IDs match those provided in the remainder of the study.

**File S5** RNAseq results. A ten-column tab-delimited text file containing all nominally-significant (*P* < 0.05) differential expression tests, with the data coming directly from the Cuffdiff output. The “GeneSymbol” and “pPos_R5” columns contain the gene name and its position relative to Release 5 of the *Drosophila melanogaster* genome. The “Sample1” and “Sample2” columns give the name of the sample, the “FPKM_1” and “FPKM_2” columns give the expression level of the gene in these samples, “log2FC” gives the fold change of expression, “TestStat” gives the test statistic, “P” gives the *P*-value, and “Q”, gives the FDR value based on the full set of tests conduction for each pairwise sample contrast.

## SUPPLEMENTAL FIGURE LEGENDS

**Figure S1** Sequence of the *Ugt86Dd* 22-bp InDel region in the 15 DSPR founder strains. The stretch of DNA is within coding exon 2 of the *Ugt86Dd* gene, and the genome coordinates are relative to Release 6 of the *Drosophila melanogaster* reference. SNPs in red are synonymous. The 22-bp deletion in founders A3, B6, B7, and AB8 leads to a short incorrect amino acid sequence (blue) followed by a premature stop codon. The sequence highlighted in yellow is the guide RNA sequence used to generate double-strand breaks (“DSB”) via CRISPR/Cas9 editing in A4.

**Figure S2** Results of quantitative complementation tests. We carried out tests using a series of deficiencies and insertional mutants. The “Interaction” *P*-value presented assesses the significance of the Founder × Mutant interaction, determining whether there is a significant quantitative failure to complement. Information regarding the positions of deficiencies/insertions is reported based on Release 6 of the *Drosophila melanogaster* reference genome. Information regarding the genes deleted was taken from FlyBase on March 11, 2017.

## SUPPLEMENTAL TABLE INFORMATION

**Table S1** Phenotypes of multiple *Ugt86Dd* CRISPR/Cas9-derived homozygous third chromosome genotypes.

**Table S2** The estimated frequency of the insertion and deletion alleles at *Ugt86Dd* in various natural populations.

